# Blood vessel occlusion by *Cryptococcus neoformans* is a mechanism for haemorrhagic dissemination of infection

**DOI:** 10.1101/2020.03.17.995571

**Authors:** Josie F Gibson, Aleksandra Bojarczuk, Robert J Evans, Alfred Kamuyango, Richard Hotham, Anne K Lagendijk, Benjamin M Hogan, Philip W Ingham, Stephen A Renshaw, Simon A Johnston

## Abstract

Meningitis caused by infectious pathogens are associated with vessel damage and infarct formation, however the physiological cause is unknown. *Cryptococcus neoformans*, is a human fungal pathogen and causative agent of cryptococcal meningitis, where vascular events are observed in up to 30% of cases, predominantly in severe infection. Therefore, we aimed to investigate how infection may lead to vessel damage and associated pathogen dissemination using a zebrafish model for *in vivo* live imaging. We find that cryptococcal cells become trapped within the vasculature (dependent on there size) and proliferate there resulting in vasodilation. Localised cryptococcal growth, originating from a single or small number of cryptococcal cells in the vasculature was associated with sites of dissemination and simultaneously with loss of blood vessel integrity. Using a cell-cell junction tension reporter we identified dissemination from intact blood vessels and where vessel rupture occurred. Finally, we manipulated blood vessel stifness via cell junctions and found increased stiffness resulted in increased dissemination. Therefore, global vascular vasodilation occurs following infection, resulting in increased vessel tension which subsequently increases dissemination events, representing a positive feedback loop. Thus, we identify a mechanism for blood vessel damage during cryptococcal infection that may represent a cause of vascular damage and cortical infarction more generally in infective meningitis.

## Introduction

Life threatening systemic infection commonly results from tissue invasion requiring dissemination of microbes, usually via the blood stream. Blood vessel damage and blockage are commonly associated with blood infection, as exemplified by mycotic (infective) aneurisms or sub-arachnoid haemorrhage (1). Indeed, both bacterial and fungal meningitis are associated with vascular events including vasculitis, aneurisms and infarcts (1–5).

The mechanisms of dissemination to the brain in meningitis have been extensively studied *in vitro* and *in vivo*. Experimental studies suggest three potential mechanisms: passage of the pathogen between cells of the blood brain barrier, polarised endocytosis and exocytosis of the pathogen by brain vascular endothelial cells, and passage through the blood brain barrier inside immune cells. However, we hypothesised that blood vessel blockage and haemorrhagic dissemination might be an alternative mechanism.

*Cryptococcus neoformans* is an opportunistic fungal pathogen causing life threatening cryptococcal meningitis in severely immunocompromised patients. *C. neoformans* is a significant pathogen of HIV/AIDs positive individuals with cryptococcal meningitis ultimately responsible for 15% of all AIDS related deaths worldwide (6). *C*. *neoformans* has previsously been suggested to disseminate from the blood stream into the brain through different routes, including transcytosis, and by using phagocytes as a Trojan horse (7–11). However, in support of our hypothesis a small number of clinical studies have suggested that blood vessel damage and bursting may also facilitate cryptococcal dissemination. Case reports indicate that cortical infarcts are secondary to cryptococcal meningitis, and suggest a mechanism whereby resulting inflammation may cause damage to blood vessels (12–14). In retrospective studies of human cryptococcal infection, instances of vascular events resulting in infarcts were seen in 30% of cases, predominantly within severe cases of cryptococcal meningitis (15).

There are two large challenges in understanding dissemination during infection that have limited mechanistic study. Firstly, the requirement for serial live imaging of a whole animal over hours or days. Secondly, the large variation in microbial pathogenesis and virulence including but not limited to hyphal invasion (3), haemolytic toxin production (16) and thrombosis (4). Long term *in vivo* analysis of infection is not possible in mammalian models; in zebrafish, by contrast, the ease of imaging infection enables visualisation of infection dynamics over many days (17,18). We observed cryptococcal cells becoming trapped and subsequently proliferating within the vasculature. Analysis of the dynamics of infection, via mixed infection of two fluorescent strains of *C. neoformans*, demonstrated that cryptococcomas within small blood vessels were responsible for overwhelming systemic infection. Localised expansion of *C. neoformans* was observed at sites of dissemination into surrounding tissue. Using a new VE-cadherin transgenic reporter line, we identified physical damage to the vasculature at sites of cryptococcal colonisation and found that blood vessels respond to their colonisation via expansion. Thus, our data demonstrate a previously uncharacterised mechanism of cryptococcal dissemination from the vasculature, through trapping, proliferation, localised blood vessel damage and through a global vasodilation response.

## Results

### Individual cryptococcal cells arrest in blood vessels and form masses

Infection of zebrafish with a low dose of ∼25 CFU of *C. neoformans*, directly into the bloodstream resulted in single cryptococcal cells arrested in the vasculature (Fig. 1A). We found that individual cryptococcal cells were almost exclusively trapped in the narrow inter-segmental and brain vessels, it is noteworthy that these vessels are similar in size to mouse brain blood vessels (Fig. 1B; (19,20)). Studies using intravital imaging in mice have previously noted cryptococcal cell trapping but due to the limitations of this model the effect of this phenomenon on disease could not be established. Exploiting the unique capacity of zebrafish for long term, non-invasive *in vivo* imaging, we found that the sites of single or very small numbers of trapped cells progressed to form cryptococcal masses or cryptococcomas within blood vessels (Fig. 1C). We found no evidence of cryptococcomas movement along vessels once established and occlusion by cryptococcomas was sufficient to prevent passage of blood cells in blocked vessel (Fig. 1D). Cryptococcomas imaged with a cytoplasmic GFP marker did not make direct contact with the vessel wall, due to the presence of the cryptococcal polysaccharide capsule, visualised by antibody staining, which also enveloped large cryptococcal masses (Fig. 1E-G). Thus, we could demonstrate that single cryptococcal cells were trapped in blood vessels and appeared to proliferate to form cryptococcal masses encased in polysaccharide capsule.

**Figure 1.**
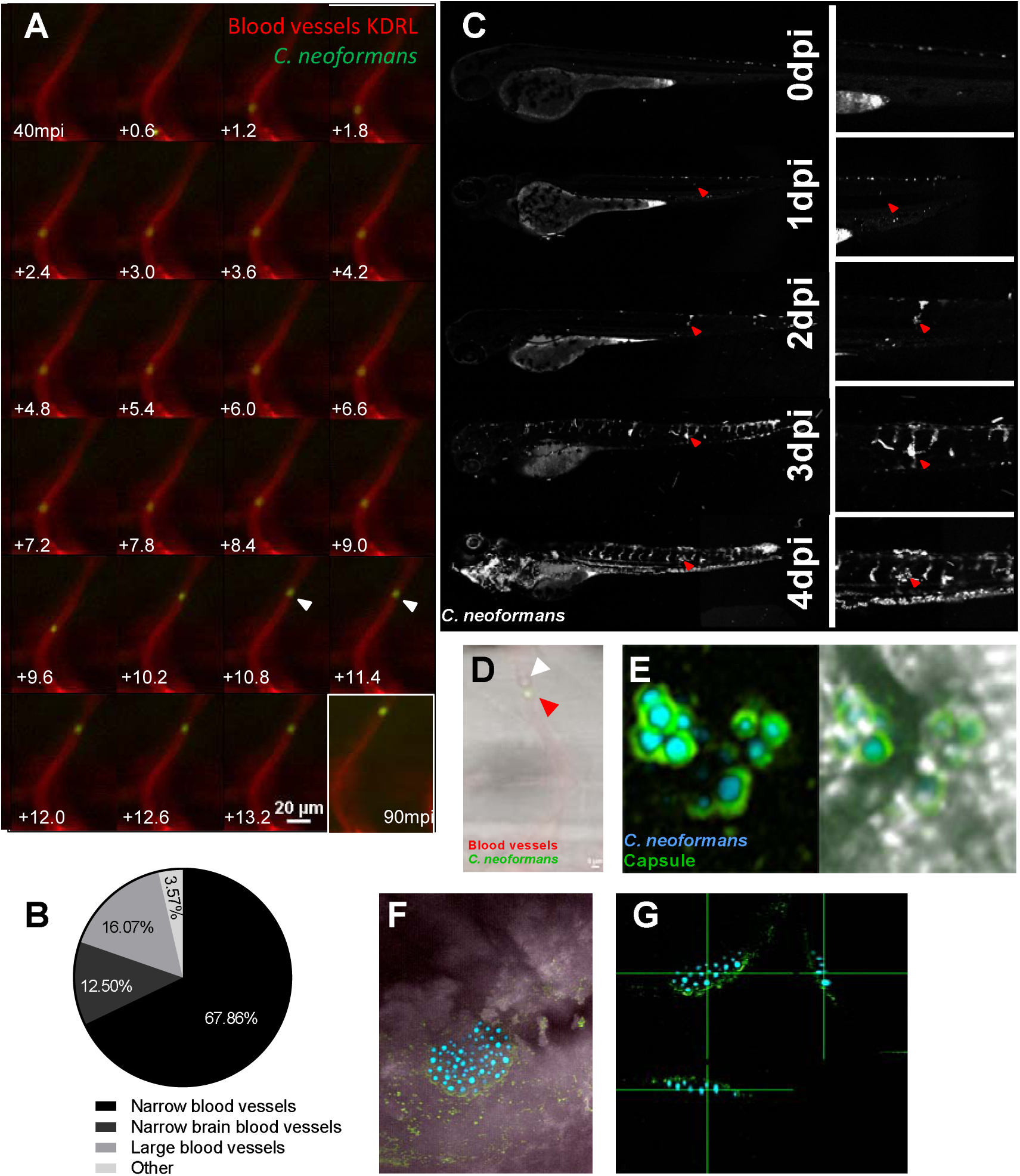
Cryptococcoma formation by cryptococcal cell trapping in small blood vessels in the zebrafish. **A** Infection of KDRL mCherry blood marker transgenic line with 25cfu GFP *C. neoformans*, imaged immediately after infection. A single cryptococcal cell becomes trapped in the vasculature (white arrow), at 40 minutes post infection (mpi) after moving from the bottom of the vessel toward the top (left to right, time points +0.6 seconds). Last image shows cryptococcal cell in the same location at the end of the timelapse at 90mpi **B** Infection of 2dpf AB larvae with 25cfu of a 5:1 ratio of GFP:mCherry KN99 *C. neoformans*. Larvae were imaged until 8dpf, or death (n=3, in each repeat 7, 10 and 12 larvae were used) Proportion of cryptococcomas observed in small inter-somal blood vessels, small brain blood vessels, large caudal vein or in other locations e.g. yolk, (n=3). **C** Infection of 2dpf AB larvae with 25cfu of a 5:1 ratio of GFP:mCherry KN99 *C. neoformans*. Larvae were imaged until 8dpf, or death (n=3, in each repeat 7, 10 and 12 larvae were used). In this case an mCherry majority overwhelming infection was reached. Infection progression from 0dpi (day of infection imaged 2hpi), until 4dpi. Red arrows follows an individual cryptococcoma formation and ultimate dissemination. **D** Infection of 2dpf AB larvae with 1000/25cfu of a 5:1 ratio of GFP:mCherry KN99 *C. neoformans* showing blood cells trapped behind a cryptococcal mass within an inter-segmental vessel. **E-G** GFP KN99 (cyan), antibody labelled cryptococcal capsule (green).**E** Cryptococci within blood vessels demonstrating the enlarged capsule blocking the vessel 24 hpi **F-G** Cryptococcal mass encased in capsule. **F**. Merged florescence and transmitted light z projection **G** Three-dimensional section of cryptococcal mass showing encasement in polysaccharide capsule.

### Clonal expansion of cryptococcal in small vessels results is associated with high fungal burden

Examination of infection dynamics over time revealed that cryptoccocal masses were often present before overwhelming infection and death (Fig. 2A). This suggested to us that cryptococcoma formation might represent a population “bottle-neck”. Several bacterial pathogens have been demonstrated to establish disease via a population “bottle-neck” i.e. clonal expansion of an individual or small number of pathogens (21,22). Initially, we injected a 1:1 ratio of GFP and mCherry-labelled cryptococci and found that single colour infections were very rare. Therefore, we decided to use a skewed ratio so that we could better quantify the likelihood of a population “bottleneck” during the progression of cryptococcal infection. We injected 25cfu of a 5:1 ratio of GFP and mCherry-labelled cryptococci and followed the infections for up to 7days post infection (dpi). In 51.6% of all infected larvae, a high fungal burden end-point was demonstrated with either cryptococci observed in the larvae were predominantly GFP positive, predominantly mCherry positive cryptococci, or a mixed outcome of both GFP and mCherry positive cryptococci (Fig. 2B). In the remaining 48.4% of infected larvae were overwhelmed by infection, or were able to clear infection so were not included in this study. Interestingly, a mixed final outcome group was not a rare occurrence (Fig. 2C). The high proportion of mixed GFP and mCherry overwhelming infections demonstrated that a single cryptococcal cell was highly unlikely to give rise to the final infection population. The predominantly GFP positive outcome group was observed most often, but only for 56.25% of all endpoints. This was far lower than would be expected, given the initial 5:1 ratio of differently labelled cells injected. While a 5:1 ratio of GFP:mCherry was injected into each larva, the actual number and ratio of cryptococcal cells varied between individual fish (Fig. 2D, SFig 1). When single colour and mixed outcomes where compared there was no significant difference in the injected ratio (Fig. 2E). However, correlative analysis demonstrated that there was no relationship between the initial ratio and final outcome ratio, suggesting there were occurrences of clonal expansion during infection (Fig. 2F). Therefore, it appeared that, while a population “bottleneck” was not common in the progression of uncontrolled cryptococcal infection, there was a skewing in the cryptococci that contributed to the final population. We hypothesized that this skewing was determined by the clonal expansion of cryptococcal masses within blood vessels. To test this hypothesis, we analysed the colours of cryptococcal masses following infection with a 1:1 ratio of GFP and mCherry-labelled cryptococci and we found that masses were of a single colour in 14/15 infections 3 dpi. This suggests that skewing of the cryptococcal population within the fish occurs at the cryptococcoma stage of infection before the final infection outcome.

**Figure 2.**
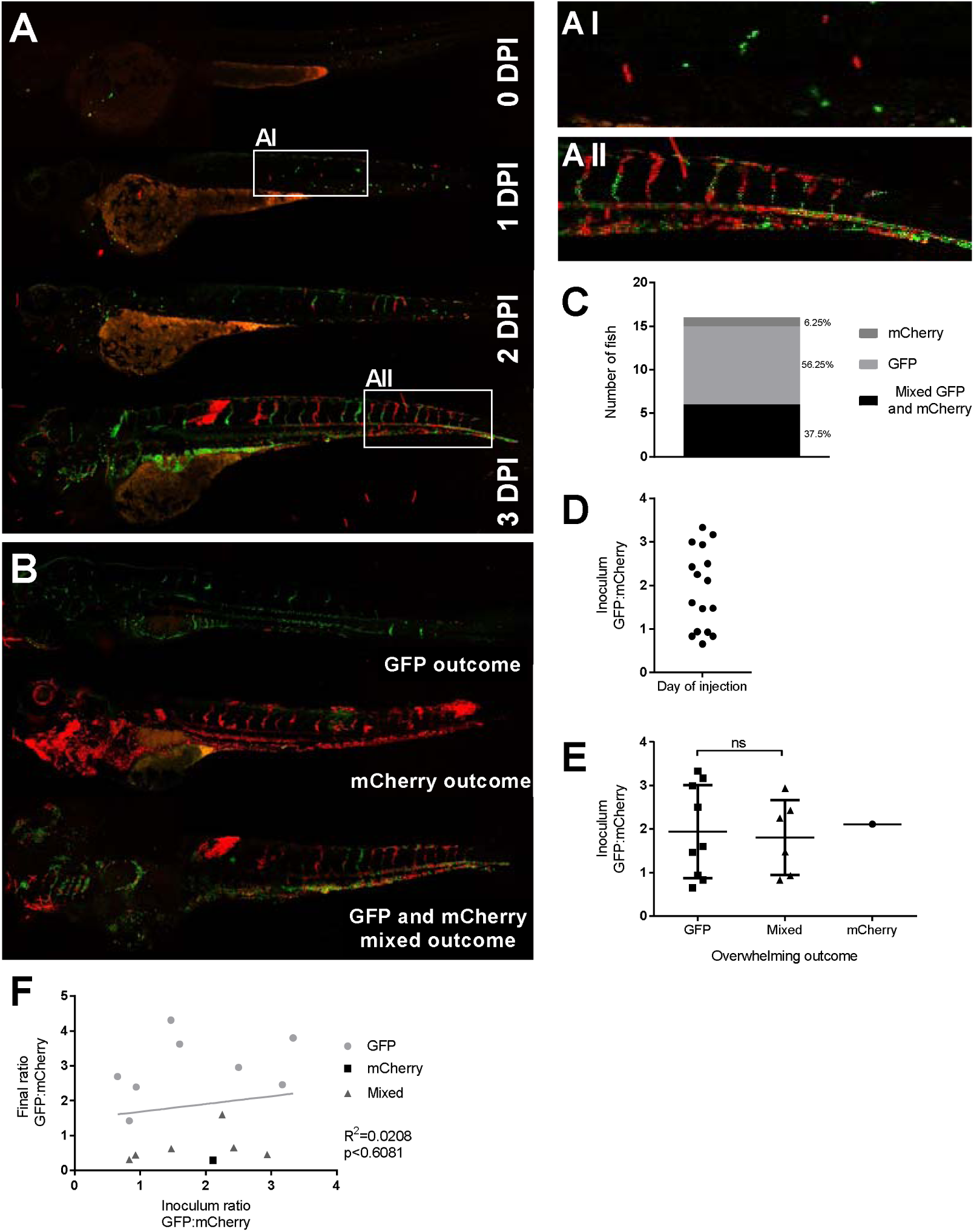
Inoculum does not predict infection outcome. Infection of 2dpf AB larvae with 25cfu of a 5:1 ratio of GFP:mCherry KN99 *C. neoformans*. Larvae were imaged until 8dpf, or death (n=3, in each repeat 7, 10 and 12 larvae were used) **A** Infection of AB wild-type larvae with 5:1 ratio of GFP:mCherry KN99 *C. neoformans*, at 0dpi, 1dpi, 2dpi and 3dpi **A I** Formation of cryptococcal masses at 1dpi **A II** Final infection outcome **B** Infection of 2dpf AB larvae with 25cfu of a 5:1 ratio of GFP:mCherry KN99 *C. neoformans*. Larvae were imaged until 8dpf, or death (n=3, in each repeat 7, 10 and 12 larvae were used). A GFP majority infection outcome, mCherry infection outcome or a Mixed GFP and mCherry infection outcome (n=3, 16 larvae) **C** Proportion of each overwhelming infection outcome observed, GFP, mCherry or mixed **D** Range of GFP:mChery *C. neoformans* injected into larvae at 2hpi **E** Actual injected GFP:mCherry ratios for each overwhelming outcome (n=3, +/- SEM, Man-Whitney t-test ns=not significant) **F** Inoculum ratio of GFP:mCherry, against final GFP:mCherry ratio at overwhelming infection stage (Linear regression R^2^=0.0208, p<0.6081, n=3, 16 larvae)

Cryptococcal masses were observed in every case preceding disseminated infection by an average of 2 days (Fig. 3A) and the number of cryptococcal masses was correlated with the rate of infection progression (Fig. 3B). We had found that individual cryptococcal cells became trapped in the narrow inter-segmental vessels (ISVs) and brain vessels, similar in size to those identified in blood vessels in the mouse brain (Fig. 1A). We quantified the distribution of cryptococcomas and found that most (80.3%) were located in these smaller brain and inter-segmental blood vessels (Fig. 1B). As cryptococcal mass formation at the start of infection was observed in the smaller blood vessels, we determined whether clonal expansion was favoured in smaller blood vessels later in infection. We compared the ratio of GFP:mCherry between the trunk blood vessels and the caudal vein and found that in mixed infections there were single colour masses in the trunk vessels but dual colours in the larger caudal vein (Fig. 2A; Fig. 3C) suggesting cryptococcal expansion occurs at sites of trapping in narrow blood vessels. Finally, to establish a role of cryptococcal masses in determining the population of cryptococci that contributed to high fungal burden, we compared the colours of individual cryptococcal masses with majority colour of high fungal burdens within individual fish. A clear relationship was demonstrated between each colour of cryptococcal masses and the disseminated infection; a single (GFP or mCherry) cryptococcal mass colour was significantly more likely to result in a single colour final outcome, with a corresponding finding for mixed cryptococcomas (Fig. 3D, E p<0.01). Together these observations suggested cryptococcal cells became trapped in small blood vessels followed by localised clonal expansion and a “skewing” of the cryptococcal population.

**Figure 3.**
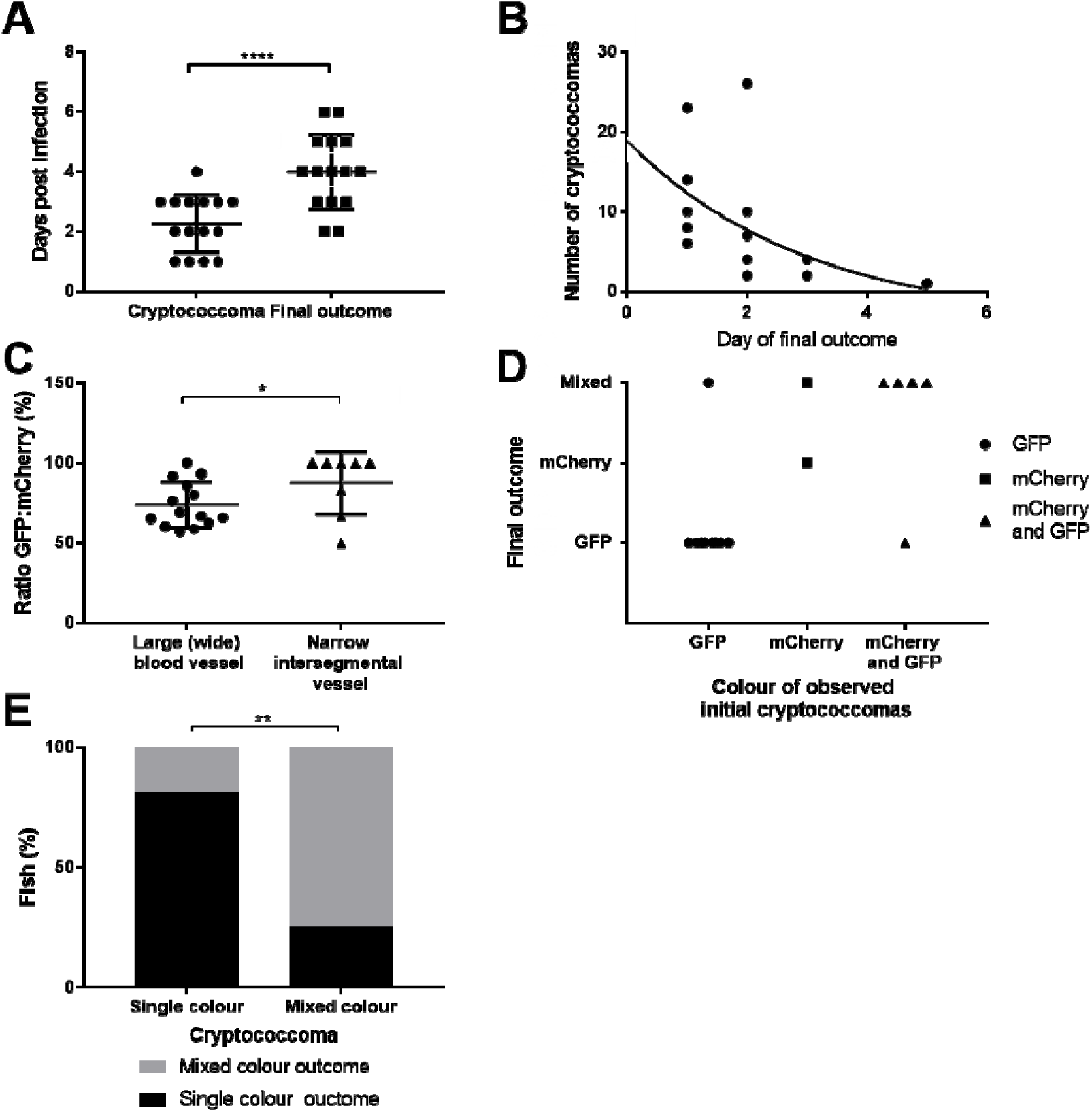
Cryptococcoma formation leads to uncontrolled infection. Infection of 2dpf AB larvae with 25cfu of a 5:1 ratio of GFP:mCherry KN99 *C. neoformans*. Larvae were imaged until 8dpf, or death (n=3, in each repeat 7, 10 and 12 larvae were used) **A** Time cryptococcoma first observed and time of final outcome observed (n=3, +/- SEM, Wilcoxon matched pairs test, ****p<0.0001) **B** The number of cryptococcomas observed within individual larvae and how many days after observation final overwhelming infection was reached (n=3, non-linear regression, one-phase decay) **C** The ratio of GFP:mCherry *C*.*neoformans* in the large caudal vein in comparison to the fifth inter-somal blood vessel, at uncontrolled infection time point (n=3, *p<0.05, +/- SEM, paired t-test). **D** Comparison of the colour (either GFP, mCherry or mixed) of *C. neoformans* in cryptococcomas, in relation to the final outcome majority *C. neoformans* colour **E** Comparison of the colour of cryptococcomas, either single colour or mixed, with the colour of final outcome (n=3, **p<0.01, Fischer’s exact test)

### Cryptococcal masses cause local and peripheral vasodilation

The finding that cryptococcal masses blocked blood vessels prompted us to measure blood vessel width at sites with or without cryptococcomas. We found that blood vessels that contained cryptococcal cells were significantly wider than those devoid of cryptococcal cells in the same infections (Fig 4A, B). A higher infection dose was used to increase the number of cryptococcal masses that formed. There was a significant difference very early, at 2 hours post infection (hpi), and a much larger difference at 3dpi (Fig. 4A, B), suggesting to us an immediate passive physical effect (i.e. due to the elasticity of the vessel wall) and a slower physical widening of the vessel caused by cryptococcal growth. We tested the first hypothesis of a fast response of the blood vessel by live imaging small brain vessels and observed that vessels locally dilated shortly after blockages formed (Fig 4C). The increase in vessel width was proportional to the size of the cryptococcal mass inside the vessel at both 2hpi and 3dpi (Fig. 4D, E) suggesting a slow increase in vessel width due to growth of the cryptococcal mass pushing against the vessel wall. Injection of inert beads of a corresponding average cryptococcal cell size (4.5μm) did not lead to formation of large masses, although there was a small but significant increase in vessel size at locations where beads did become trapped in the vasculature by 3dpi (Fig. 4F). Additionally, beads were observed stuck in the inter-segmental blood vessels much less frequently than live cryptococcal cells, with 13.6% of blood vessels containing beads compared to 89.0% containing cryptococcal cells. In addition, we specifically imaged the small vessels of the brain and found that infected blood vessels were larger relative to blood vessels in the same location in control animals (Fig. 4G, H). Thus, it appeared that blockage by cryptococcal cells and masses increased vessel diameter due to active and passive changes in blood vessels to reduce the total peripheral resistance. This appears similar to the role of increased peripheral resistance and vessel tension in higher frequencies of aneurysm (23)

**Figure 4.**
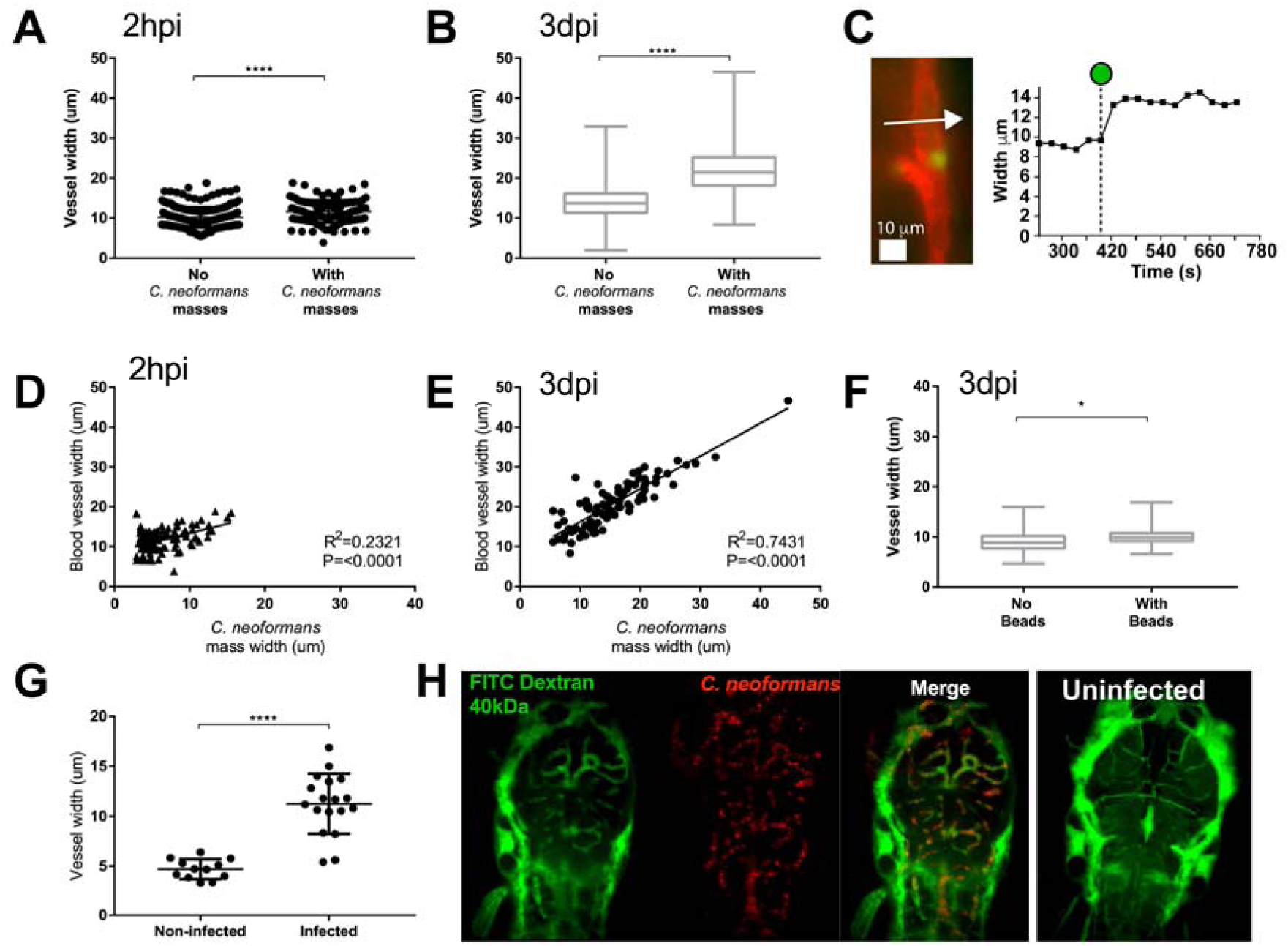
Localised clonal expansion proportionally increases vasculature size. **A-E** :Infection of KDRL mCherry blood marker transgenic line with 1000cfu GFP *C. neoformans* or inert beads **A** Vessel width with and without cryptococcal masses at 2hpi (n=3, +/- SEM, ****p<0.0001, unpaired t-test) **B** Vessel width with and without cryptococcal masses at 3dpi (n=3, +/- SEM, ****p<0.0001, unpaired t-test) **C** Left panel - Image from a time lapse movie of KDRL mCherry zebrafish larvae showing a blood vessel (red) in the zebrafish brain and a *C. neoformans* cell (green). Right panel - Kymograph showing the change in diameter of the blood vessel measured at the point indicated by the white arrow in Ci, at each frame in the time lapse. The dotted line on the x axis indicates the timepoint where the *Cryptococcus* cell becomes stuck at the point of measurement (white arrow). **D** Relationship between *C. neoformans* mass and vessel width at 2hpi (n=3, linear regression) **E** Relationship between *C. neoformans* mass and vessel width at 3dpi (n=3, linear regression) **F** Vessel width with and without beads present at 3dpi (n=3, +/- SEM, *p<0.05, unpaired t-test). **G-H**: Inoculation of mCherry *C. neoformans* with 40kDa FITC Dextran to mark blood vessels **G** Comparison of infected brain vessels width to non-infected corresponding brain vessels (three infected fish analysed, +/- SEM, ****p<0.0001, paired t-test) **H** Example image of infected and non-infected brain vessels.

### Increased cryptococcal cells size increase frequency of blood vessel occlusion

In order to investigate whether cryptococcal cell size or rigidity may affect the frequency of trapping and the extent of blood vessel vasodilation, we used mutant cryptococci with altered physical properties. Recently, the biophysical properties of several ceramide pathway mutants have been described (24) in which the accumulation of saturated GluCer (*Δsld8*) was suggested to increase the rigidity of cryptococcal membranes and therefore reduce their ability to traverse smaller blood vessels. In contrast, mutants in *Δgcs1* and *Δsmt1* have reduced amounts of the more rigid ceramide lipids or differences in lipid packing respectively. Therefore, we predicted that the *Δsld8* mutant would produce an increased number of blocked vessels whereas the *Δgcs1* and *Δsmt1* might produce reduced numbers of blockages. However, we found no differences in the number of blocked vessels or vessel width in either *Δgcs1, Δsmt1* or *Δsld8* compared to their reconstituted strains (SFig 2-4). Next, we asked whether fungal cell sized altered blockage and dilation of blood vessels. Deletion of *Δplb1* has previously been shown to exhibit increased cell size during infection of macrophages *in vitro* and in a mouse model of cryptococcosis (25,26). We have recently demonstrated that several phenotypes associated with *plb1* deletion were due to differences in fungal eicosanoid production, differences also present in a second cryptococcal mutant strain *lac1* (27). We first wanted to ensure enlarged cryptococcal size also occurs in the zebrafish infection model; we measured the size of *Δplb1* and *Δlac1* cryptococcal cells at 1dpi, and found that there was a significant increase in cell diameter compared to wild type, with a 100% increase in the number of cryptococci with a diameter >5µm (Fig. 5A). As human, rodent and zebrafish capillaries are close to 5µm at their smallest, we hypothesized that the increased fungal cell diameter of the *Δplb1* and *Δlac1* mutant cells would increase the number of vessels that would be blocked by cryptococci.

**Figure 5.**
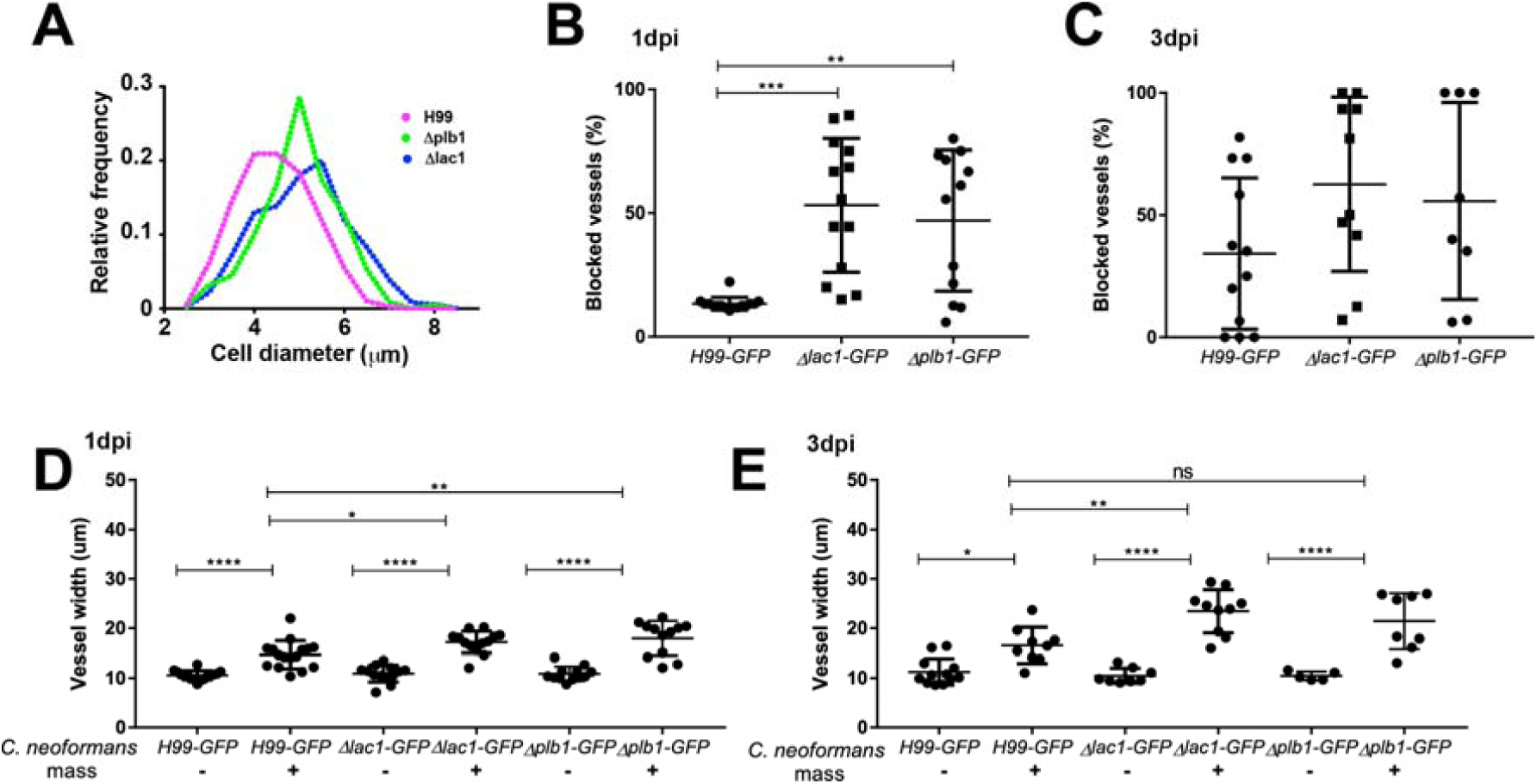
Cryptococcal cell size influences the frequency of trapping within blood vessels. **A-E:** Infection of KDRL mCherry blood marker transgenic line with 1000cfu *Δplb1-H99, Δlac1-H99* or parental *H99-GFP C. neoformans* **A** Size of cryptococcal cells injected into zebrafish larvae on the day of infection (>300 cryptococcal cells measured per strain) **B** Blocked vessels (% of inter-segmental vessels) at 1dpi (n=2, +/- SD, **p<0.01, Kruskal-Wallis test) **C** Blocked vessels (% of inter-segmental vessels) at 3dpi (n=2, +/- SD, Kruskal-Wallis test) **D** Vessel width with or without *C. neoformans* at 1dpi (n=2, +/- SD, ns=not significant, **p<0.01, ****p<0.0001, Kruskal-Wallis test) **E** Vessel width with or without *C. neoformans* at 3dpi (n=2, +/- SD, ns=not significant, *p<0.05, ****p<0.0001, Kruskal-Wallis test)

We counted the number of blocked vessels in infections with wild type, *Δplb1* and *Δlac1* mutant cryptococci and found there was a large increase in the proportion of blocked vessels at 1dpi, in some cases more than 80% of inter-segmental vessels were blocked by the *Δplb1* and *Δlac1* mutant cells (Fig. 5B). The difference in the proportion of blocked vessels was no longer significant by 3dpi (Fig 5C). We also measured the width of vessels but found no difference at either 1 or 3dpi between wild-type or mutant strains. Therefore, increased cryptococcal cell diameter led to an increase in the frequency of vessel blockages but did not significantly influence the size of cryptococcal masses, which were predominantly determined by proliferation of masses rather than individual cell size (Fig. 5D, E).

### Cryptococcal infection increases blood vessel tension resulting in hemorrhagic dissemination

Following long-term time lapse imaging of cryptococcal masses we observed that enlargement of the cryptococcoma over time eventually led to invasion of the surrounding tissue at the site of infection (Fig. 1C). The mechanism by which cryptococci disseminate from blood vessels is unknown but has been suggested to be via transcytosis or within immune cells *in vitro* (7–11). However, from our observations and from clinical reports, we hypothesised that cryptococcomas were blocking vessels, increasing the force on the blood vessel walls, leading to vessel rupture and dissemination of cryptococci. To test our hypothesis, we first established the association between tissue invasion and sites of clonal expansion within the vasculature. We found that in all cases tissue invasion occurred at sites of clonal expansion within the vasculature (19/19 tissue invasion events observed from 29 infected zebrafish; Fig 6A). Furthermore, *C. neoformans* that had invaded the surrounding tissue were invariably the same colour (GFP or mCherry) as the closest vasculature cryptococcoma (Fisher’s exact test p<0.001, n=3, Fig. 6B). To determine whether the vasculature was physically damaged sufficiently for cryptococcal cells to escape into the surrounding tissue, we examined blood vessels at high resolution at the sites of tissue invasion. We observed vessel damage and bursting at locations of cryptococcomas (Fig. 6C, D), in addition to tissue invasion events where the vasculature remained intact (Fig. 6C, 6E) but never in non-infected vessels (Fig. 6F). We could also identify possible transit by macrophages but we were unable to capture these events at sufficient time resolution to be conclusive. Blood vessel integrity is maintained by individual cell integrity and the cell-cell junctions between vascular endothelial cells. To investigate vessel integrity, through visualisation of fluorescent vessel markers, we used a stable cross of two zebrafish transgenic lines *Tg*(*10xUAS:Teal*)*uq13bh* and the endothelial *TgBAC*(*ve-cad:GALFF* (28) driver to fluorescently labell vascular endothelial cell junctional protein VE cadherin (23) in addition to the blood vessel reporter line. Using this transgenic, we also found that cryptococcal cells were located outside the blood vessel when vessels were either intact (Fig. 6E) or disrupted (Fig. 6G), in comparison to non-infected vessels (Fig. 6H).

**Figure 6.**
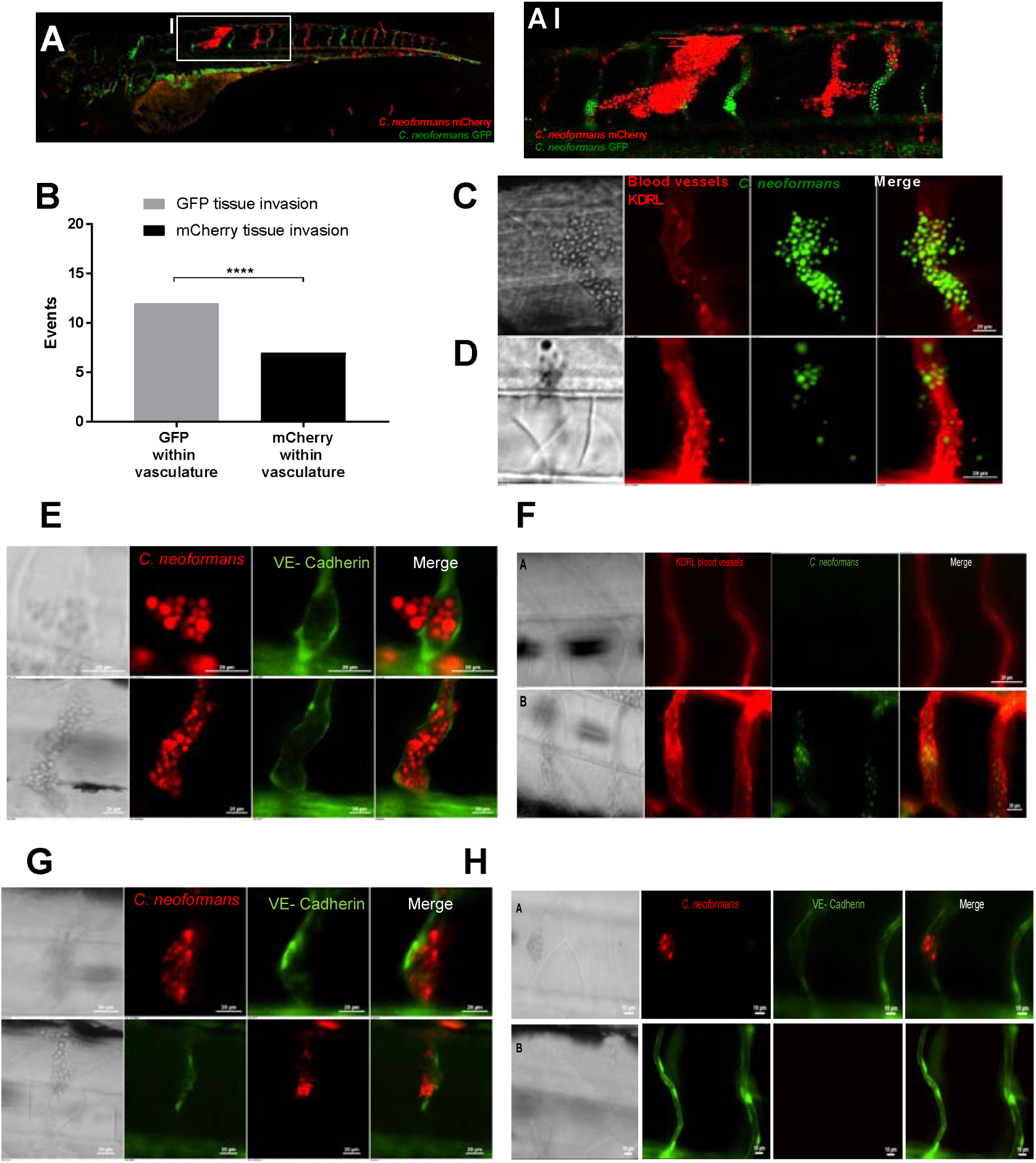
Dissemination events through vasculature damage. **A-B** Infection of 2dpf AB larvae with 25cfu of a 5:1 ratio of GFP:mCherry KN99 *C. neoformans*. Larvae were serieally imaged until 8dpf, or death **A-AI** Example of dissemination of *C. neoformans* (mCherry) into the somite surrounding an existing mCherry cryptococcoma **B** Comparison of colour of *C. neoformans* in the vasculature (GFP or mCherry), and the corresponding colour of dissemination events at the same location **C, D and F** Infection of KDRL mCherry blood marker transgenic line at 2dpf with 1000cfu GFP *C. neoformans* **C**Dissemination from an intact blood vessel, with *C. neoformans* in the surrounding tissue suggested to be transcytosis **D** Damaged blood vessels with *C. neoformans* in surrounding tissue **E**,**G and H** Infection of vascular-endothelium cadherin GFP tight junction (blood vessel marker) transgenic line with 1000cfu mCherry *C. neoformans* **E** Intact tight junctions in the blood vessel endothelial layer, with *C. neoformans* in the surrounding tissue **F** Intact blood vessels (KDRL marker) with or without *C. neoformans* **G** Damaged tight junctions in the blood vessel endothelial layer **H** Intact blood vessels (KDRL marker) with or without *C. neoformans*

We measured VE-cadherin intra-molecular tension at cell-cell junctions between vascular endothelial cells using our FRET reporter, the zebrafish transgenic line *TgBAC*(*ve-cad:ve-cadTS*)*uq11bh* (hereafter VE-cadherin-TS) (23) and found a clear decrease in VE-cadherin expression (Fig. 7A-C). In addition, we found that the VE-cadherin expression at junctions were decreased in both vessels with cryptococcal masses and those without, supporting our previous data demonstrating a global vasodilation in response to increased peripheral resistance (Fig. 7C). These, data suggested that there was an increase in vessel tension associated with cryptococcal growth in vessels and it has recently been shown that aneurysms have a higher chance of rupture under high vessel tension and peripheral resistance (23). Therefore, to test if the increased peripheral resistance was causing the haemorrhagic dissemination we had observed, we sought to increase peripheral resistance and vessel stiffness simultaneously by inhibiting the elasticity of blocked vessels. VE-cadherin is regulated extracellularly by the protease ADAM10 and inhibition of ADAM10 increases VE-cadherin junctions and blood vessel stiffness (29). Therefore, we used an ADAM 10 inhibitor during infection to increase peripheral resistance and vessel stiffness simultaneously by inhibition the elasticity of blocked vessels. In agreement with our prediction we found that inhibition of ADAM10 was sufficient to cause a large increase in the number of haemorrhagic dissemination events (Fig. 7D).

**Figure 7.**
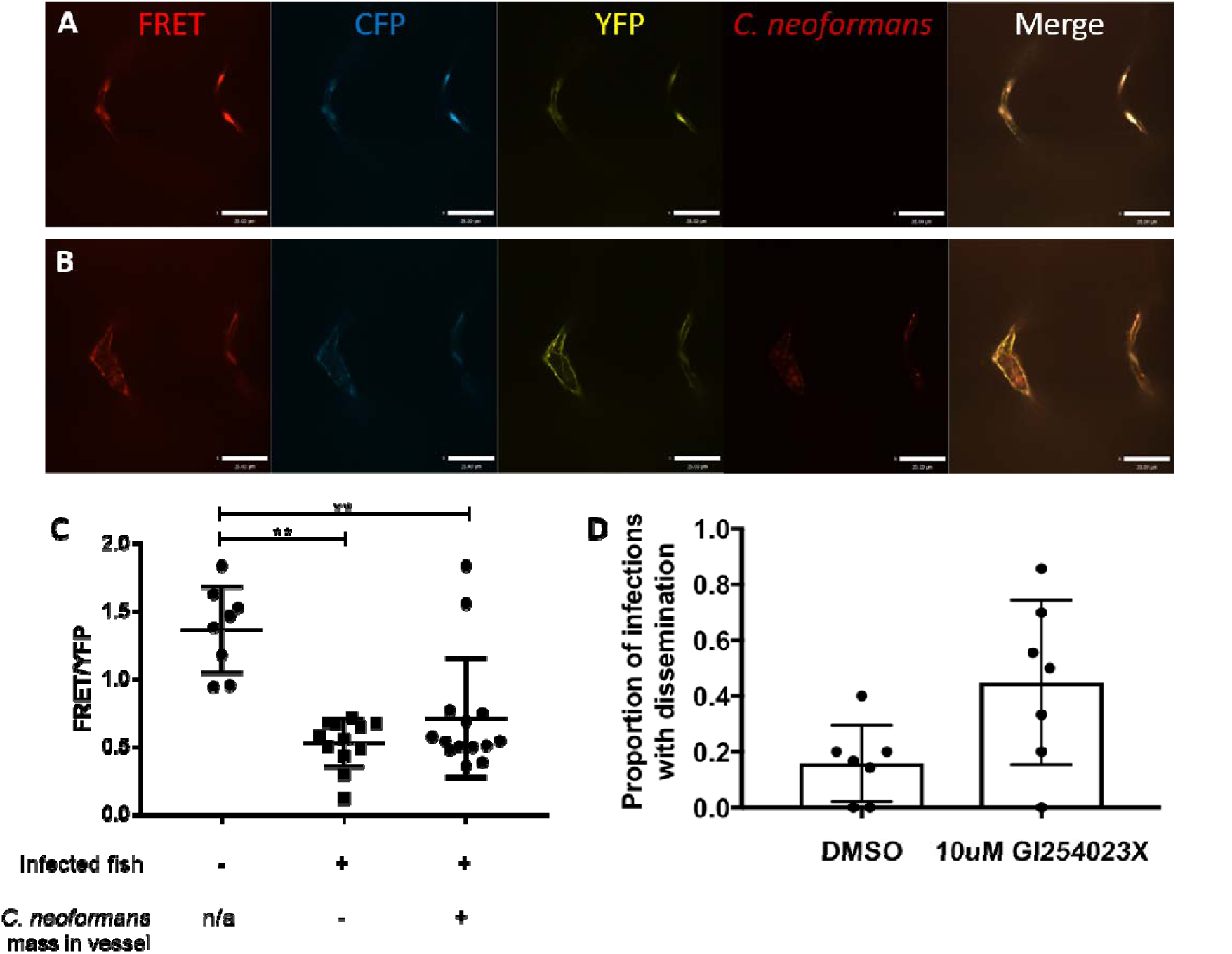
Cryptococcal infection leads to increased tension across VE-cadherin. **A-C** Infection of FRET tension reporter (VE-cadherin-TS) transgenic zebrafish line with 1000cfu mCherry *C. neoformans* **A** images of non-infected control vessels **B** Image showing infected fish, vessel containing a mass (left) and a vessel without a mass (right) **C** FRETanalysis of infected fish with or without masses and non-infected controls larvae (n=2, 4-7 larvae per repeat, +/- SD, **p<0.01, Kruskal-Wallis test, where vessel fluorescence was measured at each side of vessel) **D** Proportion of infected fish with disseminated infection. 7 repeats, 10 zebrafish larvae per repeat per group. P=0.036 unpaired t-test.

## Discussion

Here we have demonstrated how *C. neoformans* can cause haemorrhagic dissemination from blood vessels, suggesting a generallised mechanism for infarct formation during infective meningitis. Consistent with post-mortem reports showing pathogens in the brain located next to capillaries with fungal or bacterial masses present (30), our data demonstrate that, even at very low levels of fungemia, cryptococci can form masses in blood vessels, leading to increased vessel tension and blood vessel haemorrhage.

We show both localised vessel and a global vasodilation response during infection. Localised vessel vasodilation and associated damage is caused by pathogen proliferation; however the global response is likely caused by an increase in the peripheral resistance. We demonstrate that proliferation of trapped fungal cells leads to skewing of the fungal population. Importantly, we demonstrate vessel damage occurs at sites of cryptococcal masses, which likely leads to cryptococcal escape to the surrounding tissue. However, we observed dissemination via transcytosis (demonstrated where vessel structure was still intact) at sites of cryptococcomas which is suggestive that transcytosis events are also promoted by the presence of large masses of cryptococcal cells. This implicates cryptococcal growth within blood vessels in facilitating dissemination events, not only through vasculature damage.

Furthermore, we demonstrate that following cryptococcal infection a global increase in vessel vasodilation and tension across VE-cadherin occurs within the larvae, likely due to increased blood flow. We suggest increased blood flow, and therefore likely increased vessel tension, (supported with increased tension in VE-cadherin molecules) progress a positive feedback loop of increased blockages leading to further increased blood flow, vessel tension and ultimately dissemination. Indeed, we demonstrate that increasing vessel stiffness leads to increased dissemination events. This may be similar to the observed increased risk of aneurysm with high vessel tension and peripheral resistance without infection (23), perhaps further enhanced as infection progresses. Interestingly, in bacterial meningitis local damage occurs to vascular endothelial cells, but also an imbalance of hemostatic forces, potentially caused by multiple immune responses to infection, may have a systemic effect as we have shown here (31), suggesting that blood flow and vessel tension are important factors in multiple vascular infections and diseases.

The mechanism of haemorrhagic dissemination we have described for *C. neoformans* may be relevant to many infections, with multiple pathogens known to cause infarcts and vasculitis in human infection (32,33). A case report of *Candida krusei* infection in a leg ulcer causing localised vasculitis (32) suggests that local fungal pathogen growth can damage the vasculature, although this may be due to hyphal growth. In addition, infarcts are also observed in meningitis caused by bacterial pathogens, for example *S. enterica* and *T. bacillus* (34,35). Tuberculosis meningitis can also cause vasculitis leading to infarct formation (2). In bacterial meningitis, caused by *N. meningitidis*, the level of bacteraemia causes different types of vascular damage. At low bacterial numbers, bacteria are able to colonise brain blood vessels and cause limited vessel damage, eventually leading to meningitis. In contrast, high bacterial load is associated with increased vascular colonisation and augmented vascular damage (1), indicating that higher blood vessel blockage can cause increased blood vessel damage, perhaps in a similar positive feedback loop as we suggest for cryptococcal meningitis. The mechanism of vessel damage and haemorrhage may differ between species, for example *Candida albicans* infection caused haemorrhage by directly invading the blood vessel wall (5). Another aspect is the virulence factors used by the invading pathogen. In bacterial meningitis caused by virulent bacteria, *Staphylococcus aureus*, haemorrhage was observed, whereas avirulent bacteria, viridans streptococci, caused only limited blood vessel damage (16).

Vascular damage following fungal infection may not be limited to *C. neoformans, Aspergillus fumigatus* clinical reports show invasion of blood vessels could lead to hemorrhagic infarct formation (3) and in meningitis caused by *Coccidioides immitis*, infarcts were observed, at locations of thrombosis (4), potentially caused by fungal cell blockage of vessels. Furthermore, haemorrhage has been observed following an aneurysm caused by *Mucor* infection in an immuno-compromised patient with primary mucormycosis (36). Similarly, mycotic aneurysm, vasculitis and also blood vessel occlusion were observed in zygomycosis infection (37), suggesting a trapping of fungal cells and blood vessel damage may occur in different fungal species infection.

Thus, the novel mechanism of cryptococcal dissemination that we have demonstrated may be the physiological cause of infarcts observed in during blood infection. Pathogen cell trapping in narrow blood vessels, based on size, leads to localised proliferation. Growth leads to blood vessel vasodilation and damage which can allow cryptococcal cell escape into the surrounding area. In addition, cryptococcal infection induces a global vasodilation response which is associated with increased vessel tension and dissemination events. Our proposed mechanism for blood vessel bursting in cryptococcal infection may exist for other pathogens which cause vascular damage or haemorrhages, and vary depending on individual pathogen traits.

## Methods and Methods

### Ethics statement

Animal work was carried out according to guidelines and legislation set out in UK law in the Animals (Scientific Procedures) Act 1986, under Project License PPL 40/3574 or P1A4A7A5E). Ethical approval was granted by the University of Sheffield Local Ethical Review Panel. Animal work completed in Singapore was completed under the Institutional Animal Care and Use Committee (IACUC) guidelines, under the A*STAR Biological Resource Centre (BRC) approved IACUC Protocol # 140977.

### Fish husbandry

Zebrafish strains were maintained according to standard protocols (38). Animals housed in the Bateson Centre aquaria at the University of Sheffield, adult fish were maintained on a 14:10-hour light/dark cycle at 28□°C in UK Home Office approved facilities. For animals housed in IMCB, Singapore, adult fish were maintained on a 14:10-hour light/dark cycle at 28□°C in the IMCB zebrafish facility. We used the *AB* and *Nacre* strains as the wild-type larvae. The blood vessel marker *Tg(kdrl:mCherry)s916*, in addition to *Tg*(*10xUAS:Teal*)*uq13bh* (23) crossed to endothelial *TgBAC*(*ve-cad:GALFF*)(28) for stable expression. *We also used* the vascular-cadherin marker line *TgBAC(ve-cad:GALFF)* (28), *and the FRET tension sensor line, TgBAC*(*ve-cad:ve-cadTS*)*uq11bh* (23).

### *C. neoformans* culture

The *C. neoformans* variety *grubii* strain KN99, its GFP-expressing derivative KN99:GFP and mCherry-expressing derivative KN99:mCherry were used in this study (39). We used GFP expressing *Δplb1-H99, Δlac1-H99* or parental *H99-GFP* (27) *and Δgsc, Δsmt, Δsld8 and parental strain* (24). Cultures were grown in 2□ml of yeast extract peptone dextrose (YPD) (all reagents are from Sigma-Aldrich, Poole, UK unless otherwise stated) inoculated from YPD agar plates and grown for 18□hours at 28□°C, rotating horizontally at 20□rpm. Cryptococcal cells were collected from 1ml of the culture, pelleted at 3300□g for 1 minute.

To count cryptococcal cells, the pellet was re-suspended in 1□ml PBS and cells were counted with a haemocytometer. Cryptococcal cells were pelleted again (3300g) and re-suspended in autoclaved 10% Polyvinylpyrrolidinone (PVP), 0.5% Phenol Red in PBS (PVP is a polymer that increases the viscosity of the injection fluid and prevents settling of microbes in the injection needle), ready for micro-injection. The volume of PVP in Phenol red cryptococcal cells were re-suspended was calculated to give the required inoculum concertation.

### Zebrafish microinjection

An established zebrafish *C. neoformans* micro-injection protocol was followed (Bojarczuk et al., 2016). Zebrafish larvae were injected at 2 days post fertilisation (dpf) and monitored until a maximum of 10dpf. Larvae were anesthetised by immersion in 0.168□mg/mL tricaine in E3 and transferred onto 3% methyl cellulose in E3 for injection.1nl of cryptococcal cells, where 1nl contained 25cfu, 200cfu or 1000cfu, was injected into the yolk sac circulation valley. For micro-injection of GFP fluorescent beads (Fluoresbrite® YG Carboxylate Microspheres 4.50µm). The bead stock solution was pelleted at 78g for 3 minutes, and re-suspended in PVP in phenol red as above for the required concentration. Micro-injection of 40kDa FITC-dextran (Sigma-Aldrich) at 3dpf in a 50:50 dilution in PVP in phenol red, injected 1nl into the duct of Cuvier. Larvae were transferred to fresh E3 to recover from anaesthetic. Any zebrafish injured by the needle/micro-injection, or where infection was not visually confirmed with the presence of Phenol Red, were removed from the procedure. Zebrafish were maintained at 28□°C.

### Microscopy of infected zebrafish

Larvae were anaesthetized 0.168□mg/mL tricaine in E3 and mounted in 0.8% low melting agarose onto glass bottom microwell dishes (MatTek P35G-1.5-14C). For low *C. neoformans* dose infection time points, confocal imaging was completed on a Zeiss LSM700 AxioObserver, with an EC Plan-Neofluar 10x/0.30 M27 objective. Three biological repeats contained 7, 10 and 12 infected zebrafish. Larvae were imaged in three positions to cover the entire larvae (head, trunk and tail) at 2hpi, and at subsequent 24 hour intervals. After each imaging session, larvae were recovered into fresh E3 and returned to a 96-well plate.

A custom-build wide-field microscope was used for imaging transgenic zebrafish lines blood vessel integrity after infection with *C. neoformans*. Nikon Ti-E with a CFI Plan Apochromat λ 10X, N.A.0.45 objective lens, a custom built 500□μm Piezo Z-stage (Mad City Labs, Madison, WI, USA) and using Intensilight fluorescent illumination with ET/sputtered series fluorescent filters 49002 and 49008 (Chroma, Bellow Falls, VT, USA). Images were captured with Neo sCMOS, 2560□×□2160 Format, 16.6□mm x 14.0□mm Sensor Size, 6.5□μm pixel size camera (Andor, Belfast, UK) and NIS-Elements (Nikon, Richmond, UK). Settings for *Tg(kdrl:mCherry)* and *TgBAC(ve-cad:GALFF)* crossed to *Tg(10xUAS:Teal)*^*uq13bh*^ GFP, filter 49002, 50□ms exposure, gain 4; mCherry, filter 49008, 50□ms exposure, gain 4. Settings for the GFP fluorescent beads were altered for GFP alone, filter 49002, 0.5□ms exposure, gain 4. In all cases a 50um z-stack section was imaged with 5um slices. Larvae were imaged at 2hpi, and at subsequent 24 hour intervals. After each imaging session, larvae were recovered into fresh E3 and returned to a 96-well plate.

Co-injection of 40KDa FITC dextran with cryptococcal cells for imaging of vasculature in the brain was completed on 3dpf immediately after dextran injection, using a Ziess Z1 light sheet obtained using Zen software. A W-Plan-apochromat 20x/1. UV-Vis lense was used to obtain z-stack images using the 488nm and 561nm lasers and a LP560 dichroic beam splitter.

### Time-lapse microscopy of infected zebrafish

For time-lapse imaging of low *C*.*neoformans* dose infection, larvae were anaesthetised and mounted as described above, with the addition of E3 containing 0.168□mg/mL tricaine over-laid on top of the mounted *Nacre* larvae. Images were captured on the custom-build wide-field microscope (as above), with CFI Plan Apochromat λ 10X, N.A.0.45 objective lens, using the settings; GFP, filter 49002, 50□ms exposure, gain 4; mCherry, filter 49008, 50□ms exposure, gain 4. Images were acquired with no delay (∼0.6 seconds) for 1□hour, starting <2mins after infection.

### FRET microscopy and analysis

The FRET tension sensor line, *TgBAC*(*ve-cad:ve-cadTS*)*uq11bh* larvae were infected with mCherry *C. neoformans* and mounted for imaging, as above. A spinning disc confocal microscope, UltraVIEW VoX spinning disk confocal microscope (Perkin Elmer, Cambridge, UK). A 40x oil lense (UplanSApo 40x oil (NA 1.3)) was used for imaging. TxRed, exitation 561nm with 525/640nm emission filter, CFP, exitation 440nm with 485nm emission filter, YFP, exitation 514nm with 587nm emission filter, and FRET exitation 440nm with 587nm emission filter were used as well as bright field images. All were acquired using a Hamamatsu C9100-50 EM-CCD camera. Volocity software was used. Analysis of images was completed using ImageJ software. The flourecense signal intensity of the FRET, CFP and YFP channels was measured at each side of a vessel. This was completed at the location of a cryptococcal mass, or if no mass was present the middle of the vessel was measured. The FRET signal was then divided by the YPF signal, and an average was taken per vessel.

### Image analysis

Image analysis performed to measure the size of cryptococcal masses, and blood vessel width was completed using NIS elements. Fluorescence intensity of GFP and mCherry *C. neoformans* for low infection analysis was calculated using ImageJ software.

### Statistical analysis

Statistical analysis was performed as described in the results and figure legends. We used Graph Pad Prism 6-8 for statistical tests and plots.

## Author contributions

JFG, SAJ, PWI and SAR conceived of the study and designed the experiments. JFG, RJE, AB, AK, SAJ and RH performed experiments. JFG, RJE, AB, AK, RH and SAJ analysed data. AKL and BMH provided unpublished reagents and technical advice. JFG and SAJ prepared the manuscript with input from SAR and PWI. All authors commented on and edited the manuscript.

## Acknowledgments

We thank Timothy Chico (University of Sheffield, UK) and Robert Wilkinson (University of Sheffield, UK) for help and advice on vascular biology and Mike Tomlinson (University of Birmingham) for help and advice on ADAM regulation of VE-Cadherin. We thank Arturo Casadevall (Johns Hopkins University, Maryland USA) for providing the 18B7 antibody and Maurizio Del Poeta for providing *Cryptococcus* ceramide pathway mutants. JFG was supported by an award from the Singapore A*STAR Research Attachment Programme (ARAP) in partnership with the University of Sheffield. Work in the PWI lab was funded by the A*STAR Institute of Molecular and Cell Biology (IMCB) and the Lee Kong Chian School of Medicine. RJE was supported by a British Infection Association postdoctoral fellowship (https://www.britishinfection.org/). AKL was supported by a University of Queensland Postdoctoral Fellowship. BMH by an NHMRC/National Heart Foundation Career Development Fellowship (1083811). SAJ, AB, RJE, AK and RH, were supported by Medical Research Council and Department for International Development Career Development Award Fellowship MR/J009156/1 (http://www.mrc.ac.uk/). SAJ was additionally supported by a Krebs Institute Fellowship (http://krebsinstitute.group.shef.ac.uk/), and Medical Research Council Centre grant (G0700091). RJE was supported by a British Infection Association postdoctoral fellowship. AK was supported by a Wellcome Trust Strategic Award in Medical Mycology and Fungal Immunology (097377/Z/11/Z). SAR was supported by a Medical Research Council Programme Grant (MR/M004864/1). Light sheet microscopy was carried out in the Wolfson Light Microscopy Facility, supported by a BBSRC ALERT14 award for light-sheet microscopy (BB/M012522/1). We thank aquarium staff at the Bateson Centre (Sheffield) and the IMCB (Singapore) for zebrafish husbandry.

## Supplemental figures

**Figure S1.**
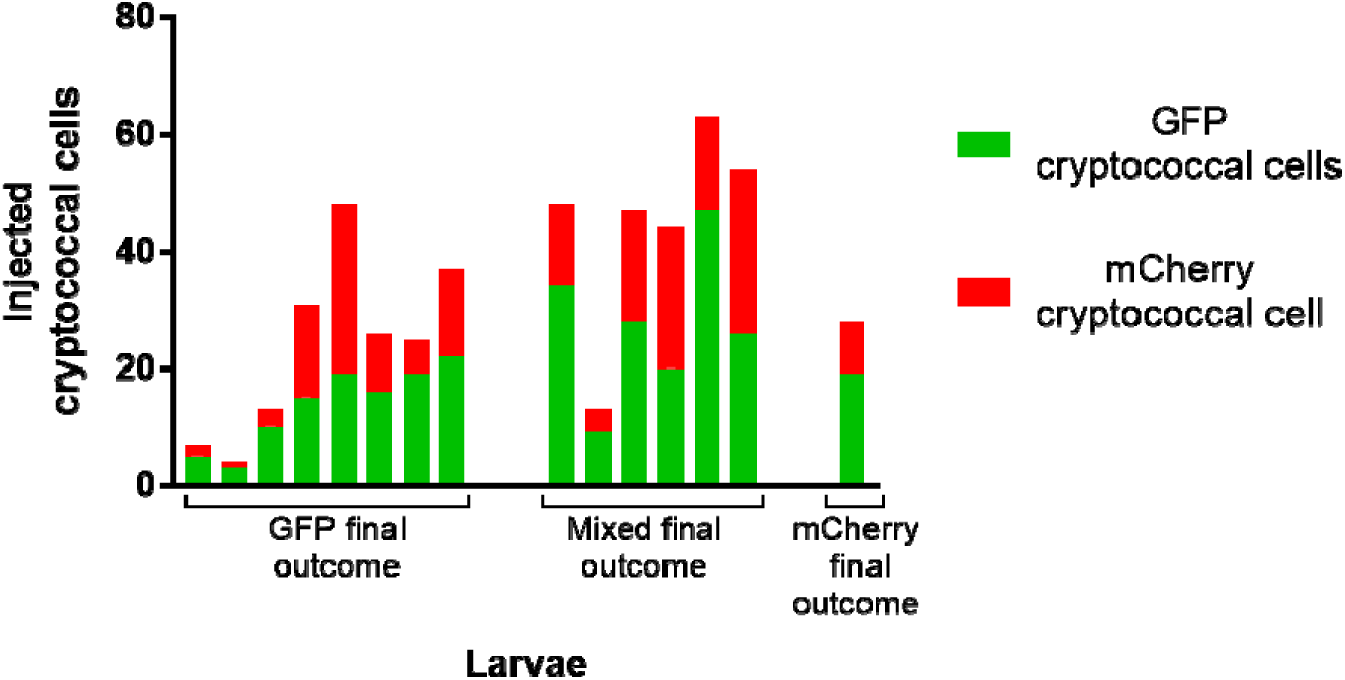
Injected ratio and number does not determine uncontrolled infection. **A** Infection of AB wild-type larvae with 5:1 ratio of GFP:mCherry KN99 *C. neoformans*, actual number of cryptococcal cells, both GFP and mCherry KN99 in 25cfu injected grouped by majority colour outcome. Each bar represents an individual fish.

**Figure S2.**
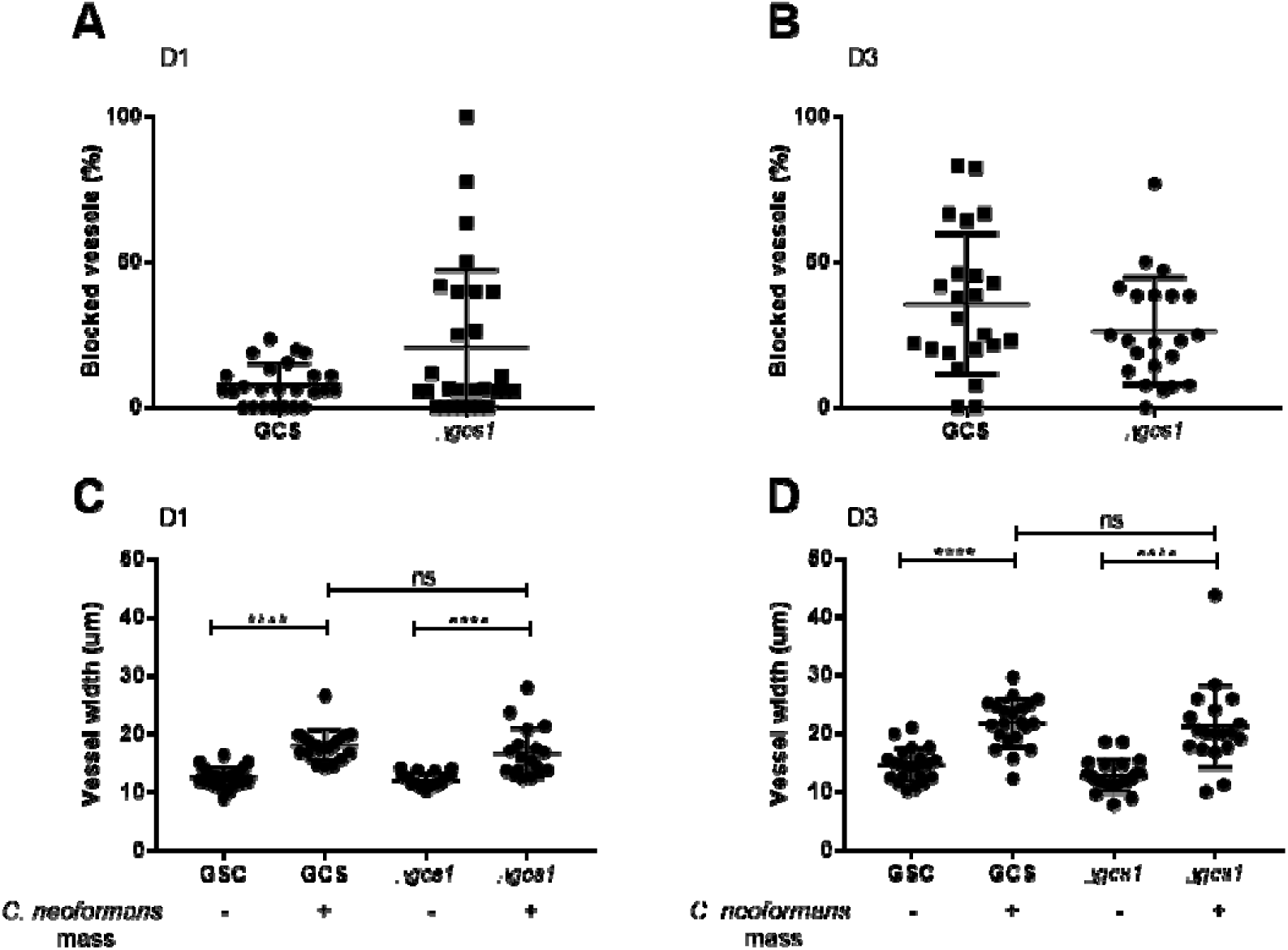
*Δgsc* does not affect blood vessel widening or frequency of trapping. **A-D:** Infection of KDRL mCherry blood marker transgenic line with 1000cfu *Δgsc* or its parental strain *C. neoformans* **A** Blocked vessels (%) at 1dpi (n=2, +/- SD, Kruskal-Wallis test) **B** Blocked vessels (%) at 3dpi (n=2, +/- SD, Kruskal-Wallis test) **C** Vessel width with or without *C. neoformans* at 1dpi (n=2, +/- SD, ns=not significant, ****p<0.0001, Kruskal-Wallis test) **D** Vessel width with or without *C. neoformans* at 3dpi (n=2, +/- SD, ns=not significant, ****p<0.0001, Kruskal-Wallis test)

**Figure S3.**
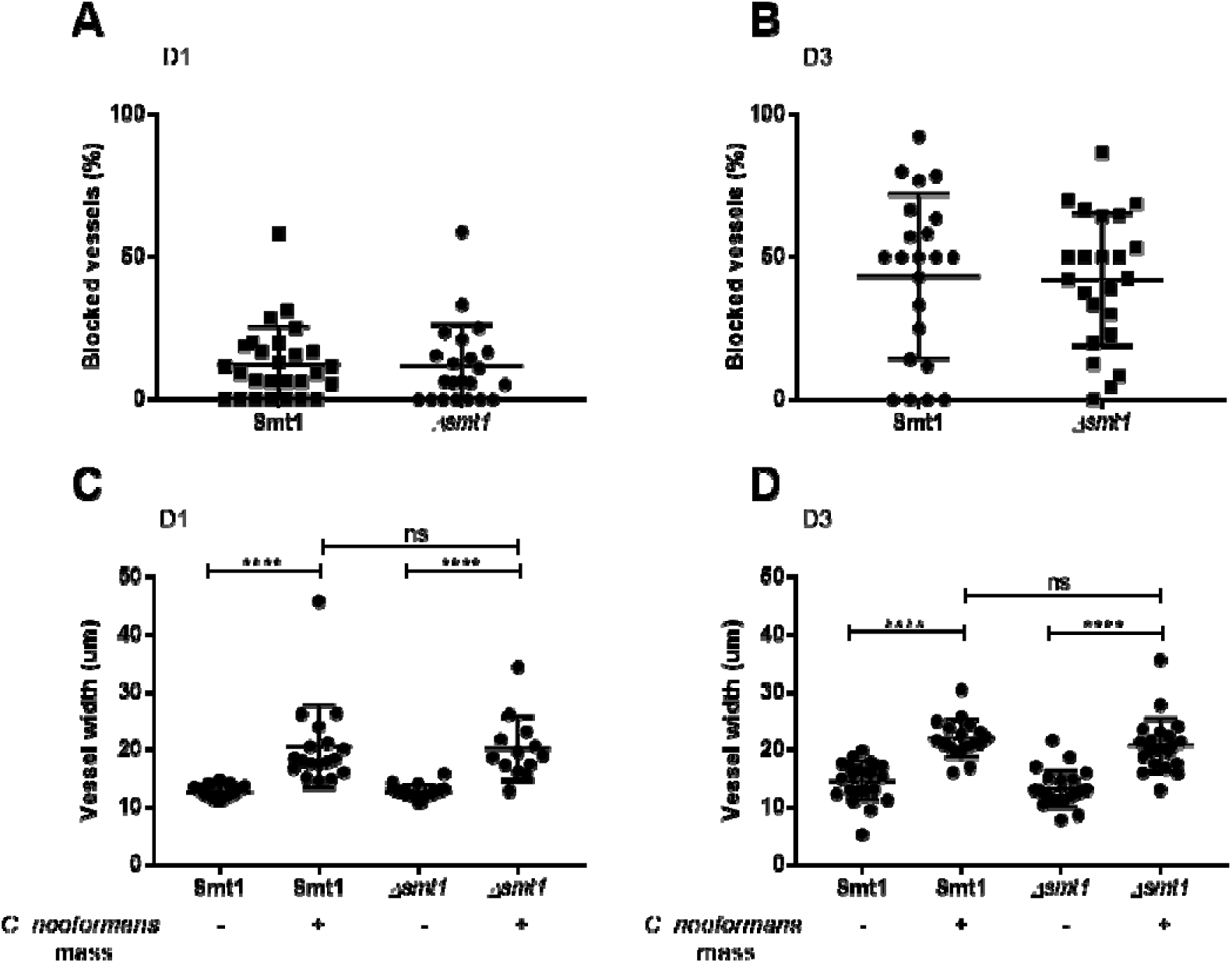
*Δsmt* does not affect blood vessel widening or frequency of trapping. **A-D:** Infection of KDRL mCherry blood marker transgenic line with 1000cfu *Δsmt* or its parental strain *C. neoformans* **A** Blocked vessels (%) at 1dpi (n=2, +/- SD, Kruskal-Wallis test) **B** Blocked vessels (%) at 3dpi (n=2, +/- SD, Kruskal-Wallis test) **C** Vessel width with or without *C. neoformans* at 1dpi (n=2, +/- SD, ns=not significant, ****p<0.0001, Kruskal-Wallis test) **D** Vessel width with or without *C. neoformans* at 3dpi (n=2, +/- SD, ns=not significant, ****p<0.0001, Kruskal-Wallis test)

**Figure S4.**
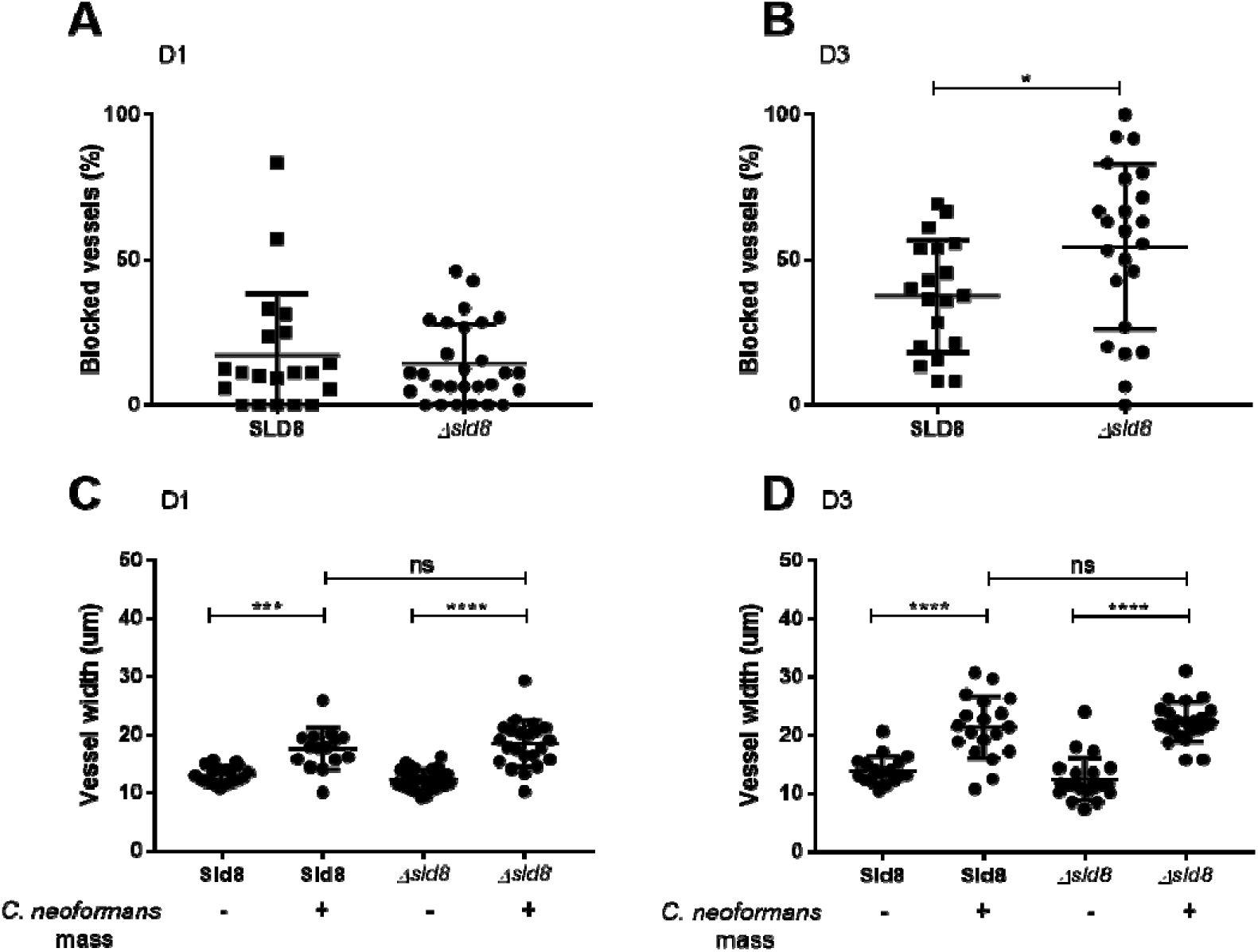
*Δsld8* does not affect blood vessel widening or frequency of trapping. **A-D:** Infection of KDRL mCherry blood marker transgenic line with 1000cfu *Δsld8* or its parental strain *C. neoformans* **A** Blocked vessels (%) at 1dpi (n=2, +/- SD, Kruskal-Wallis test) **B** Blocked vessels (%) at 3dpi (n=2, +/- SD, *p<0.05, Kruskal-Wallis test) **C** Vessel width with or without *C. neoformans* at 1dpi (n=2, +/- SD, ns=not significant, ****p<0.0001, Kruskal-Wallis test) **D** Vessel width with or without *C. neoformans* at 3dpi (n=2, +/- SD, ns=not significant, ****p<0.0001, Kruskal-Wallis test)

